# Threshold effects of prenatal stress on striatal microglia and relevant behaviors

**DOI:** 10.1101/2025.01.30.635666

**Authors:** S. V. Maurer, M. M. Evans, M. Dukle, S. Kundu, J. L. Dennis, R. M. Ellerbroek, S. L. Anema, V. C. Roshko, H. E. Stevens

## Abstract

Prenatal stress, a risk factor for neurodevelopmental disorders (NDDs), leads to immune alterations, including offspring neuroimmune cells. Differences in offspring outcomes may arise from whether the extent of prenatal stress crosses “thresholds” for effects on specific outcomes. Therefore, we sought to determine offspring outcomes using models with different extents of prenatal stress. We focused on striatal outcomes, because of their relevance for NDDs. Pregnant CD1 mice were assigned to four groups (each: N=6): no stress (“NoS”) or one of the following stressors administered three times daily: i.p. saline injections (low prenatal stress, LoS), Interleukin-6 injections as a component of prenatal stress (immune prenatal stress; ImS), or restraint stress + saline injections (high prenatal stress, HiS), embryonic day 12-18. In adult offspring, HiS altered striatal-dependent behavior across males and females, while ImS induced fewer behavioral changes, and LoS did not affect behavior. Adult striatal microglia morphologies were mostly unchanged across groups, with only HiS leading to altered striatal density of minimally ramified cells. However, embryonic striatal microglia were affected by all models of stress, albeit in distinct ways. The HiS model, and to a lesser extent LoS, also influenced immune components of the maternal-fetal interface: placental macrophages. In conclusion, high and immune stress affected adult striatal-dependent behavior, exceeding the threshold necessary for persistent impacts, but all stress models affected embryonic microglia, suggesting that early neuroimmune outcomes had a lower threshold for impacts. Distinct severities and aspects of prenatal stress may therefore underlie different outcomes relevant to NDDs.

## Introduction

Prenatal stress is a risk factor for many neurodevelopmental disorders (NDDs), including autism spectrum disorder (ASD), ADHD, and Tourette syndrome (1–5). How stress is conceptualized across human studies of prenatal stress can vary: it may be defined as a major weather event (6–12), significant life events (5, 13, 14), bereavement (2, 15–17), or through direct assessment of maternal cortisol (18–21).

Although these different stressors have associations with NDDs, there is evidence that stress experiences may have differing impacts on neurodevelopmental outcomes in children (22), demonstrating a critical need to understand the possible mechanisms behind the prenatal stress-NDD link.

Previous research in humans has suggested that the extent of stress is a factor linked to offspring neurodevelopmental outcomes (reviewed in 23). For example, severe stress is associated with an increase in neural tube defects (24), but less severe stress increases preterm birth and low birth weight risks (25, 26). As well, moderate stress due to a major weather event leads to different placental gene expression than severe stress from the same weather event (12). This suggests that there may be “thresholds” for different offspring outcomes which may depend on the extent of the stressor. Different developmental outcomes and potential intermediate mechanisms may have different “thresholds” for the impact of prenatal stress.

One potential mechanism for effects of prenatal stress is alteration of microglia, unique immune cells that reside in the central nervous system. Microglia, unlike other glial cells, arise from the yolk sac and migrate to the brain (27, 28). Because this migration happens before the full establishment of the blood-brain barrier (29), early microglia may react quickly to immune insults in the periphery and be highly responsive to external stimuli. To aid in actively maintaining homeostasis, their main roles include responding to injury, performing phagocytosis, and instigating the immune response (30–33). Thus, even small changes to microglia may have profound effects on other aspects of brain development (34, 35).

Furthermore, microglia also play a large role in the elimination of excess neural and glial cells via phagocytosis which has the potential to lead to lifelong effects and neurodevelopmental deficits (36).

Microglia are critical during development; however, few have assessed microglia in the developing striatum in the context of broad prenatal stress (37). After maternal immune activation, offspring striatal microglia have decreased immune reactivity (38). The striatum is highly relevant to NDDs (39), and striatal microglia may be critical for lasting impacts on forebrain circuitry (40).

To evaluate the thresholds for effects of prenatal stress on striatal microglia and relevant striatal-dependent behaviors, we used three models with different extents of prenatal stress. As a model with high levels of maternal stress response, we included a commonly used repetitive restraint model that has previously shown alterations to cortical microglial development and other behaviors (19, 41). We also utilized components of this full restraint stress to model different extents of stress. We defined “high” stress as repetitive restraint stress plus maternal saline injection (41), “immune” stress as repetitive maternal injection of interleukin-6 (IL-6), and “low” stress as repetitive saline injections alone. Prenatal stress leads to an increase in IL-6 in humans (42) and mice (43), and IL-6 is differentially affected by the extent of stress in humans (44). We hypothesized that “immune” and “low” stress which are components of “high” stress would show fewer alterations in striatal-dependent behaviors and striatal microglial morphology because the threshold for these effects would only be exceeded by “high” stress.

## Methods

### Stress manipulations

All animal studies were conducted with the approval of the University of Iowa IACUC committee. Wild-type CD1 females were time-mated to GAD67GFP+ CD1 male mice (45). Beginning on embryonic day 12 (E12; Figure 1A) as previously reported (41, 46–48), dams were assigned to one of 4 groups, with the specific manipulation three times per day (Figure 1*B*):

1. No stress (NoS): no handling, injections, or restraint stress
2. Low stress (LoS): saline injections
3. Immune stress (ImS): IL-6 injections, 100ng suspended in 0.2 mL sterile saline per injection
4. High stress (HiS): 0.2 mL saline injections plus full restraint stress model as previously described (41, 46–48), with dam in plexiglass restraining tube under a bright light for 45 minutes.

**Figure 1.**
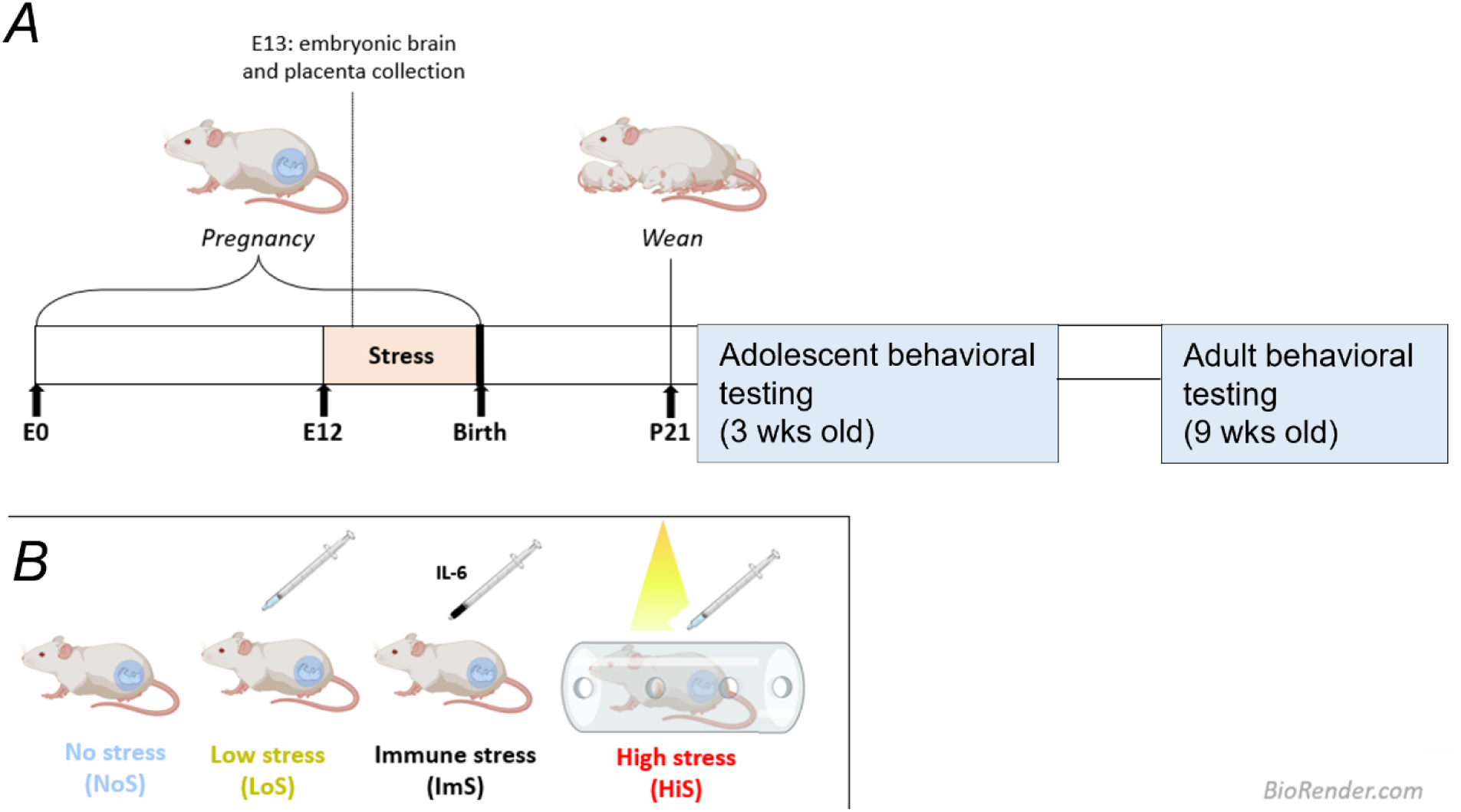
Experimental schematic. (A) Timeline of stress induction, tissue collection, and behavioral experiments. E0, E12, or E13 = embryonic day 1, 12, or 13, P21 = postnatal day 21. (B) Stress groups: No stress (NoS), Low stress (LoS), Immune stress (ImS), and High stress (HiS).

Manipulations continued until tissue collection at E13 or birth (for offspring used in behavior experiments). Offspring body weights were collected at E13, postnatal day 0 (P0), P21, and at the time of tissue collection at approximately P70.

### Behavior

Striatal-dependent behaviors were tested in independent offspring starting at P24 (adolescent) or P56 (adult). Mice underwent one testing session per day during the light cycle. Mice were habituated in a clean cage in the testing room for 30 min each day. One or two male and female littermates were tested (n = 8-14).

#### Amphetamine-induced stereotypy

Amphetamine-induced stereotypy was used to assess repetitive behaviors (49). Mice were recorded using a camera for 20 min in an arena. Then, mice were injected i.p. with 2.5 mg/kg of amphetamine, and video recordings continued. Videos were hand scored for self-circling behavior before and after amphetamine injection for one min every 5 min of the 20 min periods by a scorer blind to condition and average number/min reported.

#### Water T-maze

Water T-maze was used to assess habit learning (perseverative behavior) and cognitive flexibility in reversal training (50). The testing chamber was a pool of water, heated to about 22 degrees Celsius and with white paint for opacity, with a T-shaped plexiglass arena in the middle. The arena faced the same direction for all trials. Slightly underneath the water, a platform was placed on either the left or right side; platform location was counterbalanced across mice. Mice were placed into the bottom portion of the “T,” and timed for latency to find the platform as well as incorrect arm entries. Mice were tested for 10 trials (with 15 min intertrial intervals) per day until they reached criterion (5 correct trials in a row within one day), then trained for two more days with the platform in the same location. Reversal learning, in which the platform location was reversed, began the next day and continued until mice reached the same criterion. The number of errors (incorrect arm entries) until reaching training criterion was used to measure habit learning, and the latency on the first, second, or third reversal trial (51) was used to measure cognitive flexibility.

#### Rotarod

To assess motor learning, mice underwent a two-day rotarod task as previously described (48).

Mice underwent five trials per day, with an approximately 30 min rest between trials. “Latency” was defined as time to fall or time to second passive rotation (52). Learning rates for day one and day two were calculated as ((mean latency of trials 4 and 5) – (mean latency of trials 1 and 2))/(intertrial interval, which was 4 in these experiments)(53).

#### Open field

To assess locomotor activity, mice were placed in a plexiglass arena (40 x 39 x 30 cm) for 30 min.

They were assessed using AnyMaze (Stoelting) for time mobile and speed while mobile.

### Cellular quantification

#### Adult immunohistochemistry and stereology

Adult brain tissue at approximately P70 was perfused, post-fixed in PFA, cryoprotected in 30% sucrose, and serial cryosectioned in OCT compound from mice that were utilized for adolescent behavioral experiments. These tissues were collected at least four weeks after the completion of testing to assess the striatum for adult microglial morphology as previously described for other brain regions (41). Briefly, sections were incubated in 10% normal goat serum/PBS++ blocking solution for 1 hour.

Then, they were incubated in primary anti-IBA1 antibody (1:500, 012-26723, Wako) overnight at 4 degrees Celsius. Sections were then incubated with Alexa dye-conjugated secondary antibody (1:500; Molecular Probes), and coverslipped with a DAPI-containing mounting medium (#H-1200; Vector Laboratories). Iba1+ cells were counted in every 10^th^ 50 micron section of dorsal striatum (i.e. caudate putamen; 6-9 total sections per brain) using a AxioImager.M2 (Zeiss) coupled to Stereoinvestigator (Microbrightfield) with a counting frame of 250 x 150 by a 40x objective. Microglial morphologies were quantified as one of four states: “amoeboid” with 0-1 process; “lowly ramified” with 2-3 processes; “moderately ramified” with 4 processes *or* multiple thin, spindly processes and a small soma; or “highly ramified” with 5+ processes and a large soma (Figure 2B, 41). Density was quantified by dividing the estimated cell population by the volume of the caudate putamen.

**Figure 2.**
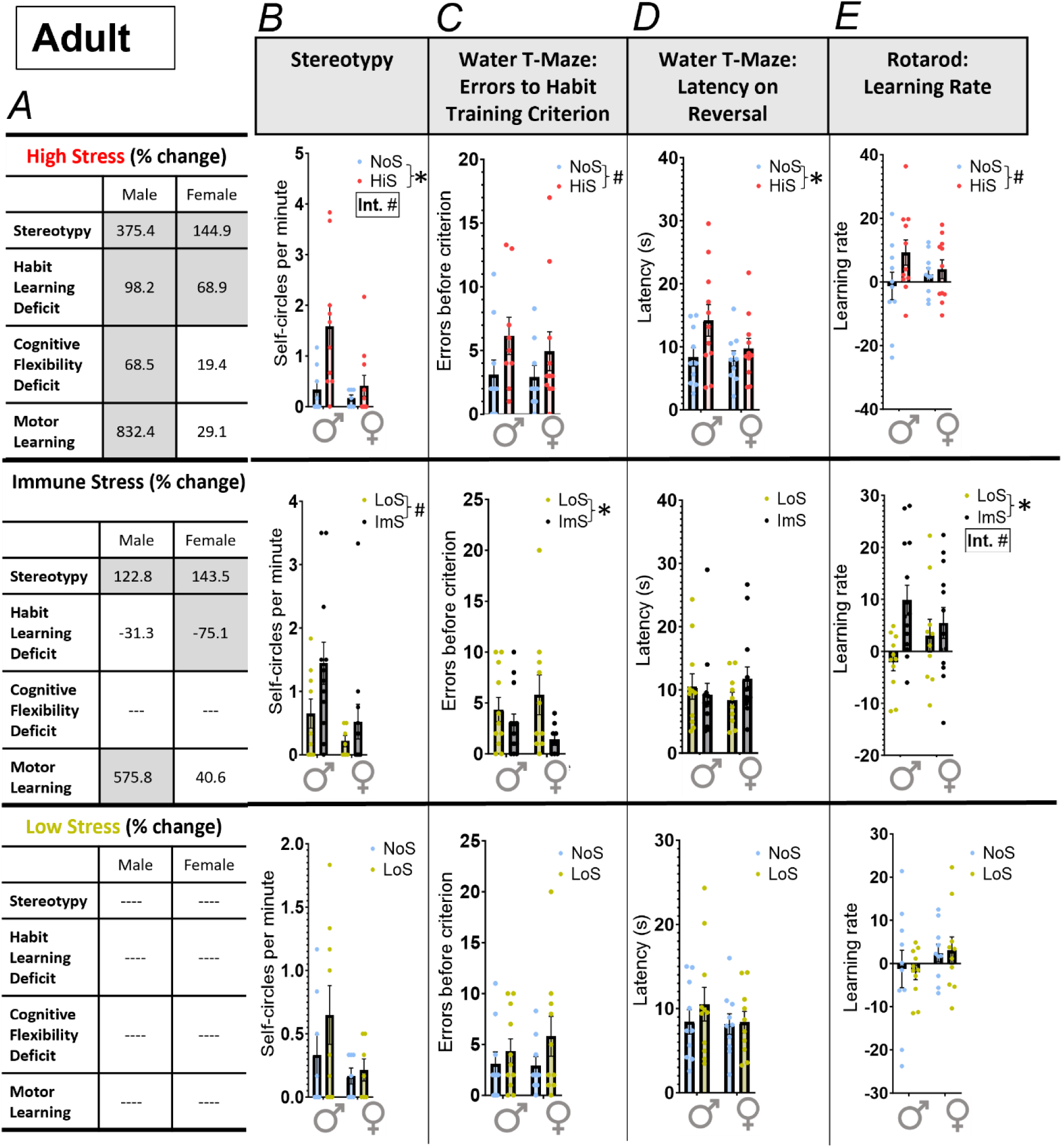
Adult striatal-dependent behaviors were largely changed by high stress in males only. * *p* < 0.05, # *p* < 0.10, “Int.” = sex x stress interaction. NoS = No stress, LoS = Low stress, ImS = Immune stress, HiS = High stress. (A) Percent changes in each task and group for outcomes with *p* < 0.10 effects. Cells that are shaded indicate a greater than 50% difference from control. (B) In adult offspring, high stress led to a significant increase in self-circling, a stereotyped behavior, after amphetamine injection. (C) Immune stress led to a decrease in errors to habit acquisition criterion, indicating an enhanced ability to acquire a new habit. High stress led to a marginal increase in errors made. (D) High-stress offspring showed a higher latency in the third reversal trial, indicating a decrease in cognitive flexibility. (E) Immune stress led to a trend and significant increase in learning rate, respectively.

#### Embryonic immunohistochemistry and cellular measures

On E13 in a separate cohort, offspring brain and placentas were collected and post-fixed in formalin. Tails were collected for sex genotyping. Brains were then cryoprotected in 30% sucrose, and cryosectioned in OCT compound at 25 microns. Placentas were microtome sectioned at 4 microns and H&E stained using the University of Iowa Comparative Pathology core. E13 brain tissue (n = 5-9) was slide-mounted, immunostained as described above (48), including additionally staining for with anti-GFP (1:1000, AB13970, Abcam) and anti-Ki67 (1:200, AB15580, Abcam) antibodies, and assessed with an Axioimager.M2.

IBA1+ cells in the E13 primordial striatum (the ganglionic eminence) across four morphologies were counted in every 20^th^ section. Cell morphologies were counted as previously described (41): “multivacuolated” with multiple nuclei and/or vacuoles present within the cell; “amoeboid” with a normal nucleus and no cellular processes; “transitional” with a normal nucleus and 1 cellular process; and “ramified” with a normal nucleus and 2+ cellular processes (Figure 5*B*). Cells were exhaustively counted (2-3 sections per brain). Density was calculated by dividing the total cell counts by the volume of the ganglionic eminence.

Migration of GAD67GFP+ cells was measured as previously described (41, 48, 54), by calculating the migratory length of the superficial stream of the dorsal forebrain cells as a percent of the length of the cortical plate (5-7 sections per brain). Ki67+ cells in the ganglionic eminence were assessed using exhaustive counting (4-5 sections per brain). The density was calculated by dividing the total cell counts by the volume of the ganglionic eminence.

After H&E staining, the labyrinth area and junctional areas of the placenta were measured using brightfield microscopy with StereoInvestigator in one section near the center of the placenta. Hofbauer cells were evaluated after antigen retrieval (boiling for 10 minutes in 10mM EDTA) and the same immunohistochemical protocol as embryonic brains was followed (1:500 anti-IBA1). Cells were exhaustively counted in the labyrinth zone, and density was calculated as the number of cells divided by labyrinth volume assessed.

### Statistical analysis

Analyses were conducted and graphs were created using GraphPad Prism v10.0.2. Outliers using the ROUT method (Q = 0.2%) were excluded. We conducted two-way ANOVAs to evaluate NoS vs HiS, LoS (control for injections) vs ImS, and NoS vs LoS comparisons. Three-way ANOVAs were utilized to assess the multiple related microglial morphologies (sex x stress x morphology). Post hoc comparisons are reported using the two-stage step-up method of Benjamini, Krieger and Yekutieli method for multiple comparisons.

## Results

### Adult striatal-dependent behavior

First, striatal-dependent behaviors were tested (Fig 2). In general, high stress exposure (HiS) changed striatal-dependent behavior the most, with immune stress exposure (ImS) having fewer but significant impacts (Fig 2*A*). Interestingly, low stress exposure (LoS) offspring were not impacted on any adult behavioral measure, indicating a higher threshold for prenatal stress effects at this timepoint for both males and females. Adult offspring showed no weight differences (Supplemental Table S1) and no effects on general locomotion in an open field (Supplemental Fig S1*A*,*B*) across the four conditions.

Specifically, HiS led to an increase in stereotyped behavior (NoS vs HiS: F_1,32_ = 7.12, *p* = 0.012) as assessed by post-amphetamine quantification of self-circles (Fig 2*B*). This effect was driven by a 375.4% increase in males, with a lower 144.9% increase in females (sex x stress interaction: F_1,32_ = 3.25, *p* = 0.08, post hoc NoS males vs HiS males *p* = 0.003, Fig 2*B*). Offspring in the ImS group had trend higher stereotyped behavior with similar 122.8% and 143.5% increases in males and females respectively (LoS vs ImS: F_1,36_ = 3.64, *p* = 0.064). All offspring groups showed similar levels of self-circling before amphetamine treatment (Supplemental Fig S1*C*).

We used the water T-maze to assess habit learning and cognitive flexibility on reversal learning (Fig 2*C*,*D*). Errors in habit learning trended higher in HiS offspring (NoS vs HiS: F_1,35_ = 3.68, *p* = 0.063, Fig 2*C*) but were significantly lower after ImS (LoS vs ImS: F_1,39_ = 5.22, *p* = 0.028), indicating opposite effects of similar magnitude. However, only HiS *in utero* impacted offspring reversal learning, again showing poorer performance of these adult offspring (NoS vs HiS: F_1,38_ = 4.17, *p* = 0.048, Figure 2*D*). This was largely driven by a 68.5% increased latency in males as testing progressed (the third reversal learning trial). All offspring groups showed similar reversal learning on the first and second trials (Supplemental Fig S1*D*,*E*).

Similar to the effects observed on stereotyped behavior, HiS and ImS both affected motor learning (Figure 2*E*). A trend increase in learning rate on the second day was found for HiS offspring (NoS vs HiS: F_1,38_ = 3.03, *p* = 0.090) with a larger increase in males. Motor learning was also strongly increased in male ImS offspring (575.8%) as shown by both a significant effect of stress and a trend sex x stress interaction (LoS vs ImS: F_1,44_ = 6.57, *p* = 0.014; sex x stress interaction: F_1,44_ = 2.86, *p* = 0.098; post hoc LoS males vs ImS males *p* = 0.004). Both of these findings indicated that stress led to an improvement in motor learning. No learning differences across were found on the first day (Supplemental Fig S1*F*).

### Adult striatal microglia

Because of the effects of HiS and ImS in offspring striatal-dependent behaviors, we next assessed the density of multiple microglial morphologies in the caudate putamen, the largest striatal structure in mice (Fig 3*A*). We categorized microglia into four morphological states which classically reflect their reactivity (Fig 3*B*). Microglia with morphological states classically associated with increased inflammation (“minimally ramified”) were sex-specifically changed in HiS offspring (NoS vs HiS: sex x stress interaction: F_1,21_ = 6.11, *p* = 0.022, Fig 3*C*), with 15.7% lower density in males and 17.7% higher density in females. Neither ImS nor LoS affected adult striatal microglial morphology, suggesting a higher threshold for these adult striatal microglial outcomes than for adult behavior. No overall changes in “ramified” microglia density (Fig 3*D*) or striatal volume were observed (Supplemental Fig S1*G*).

**Figure 3.**
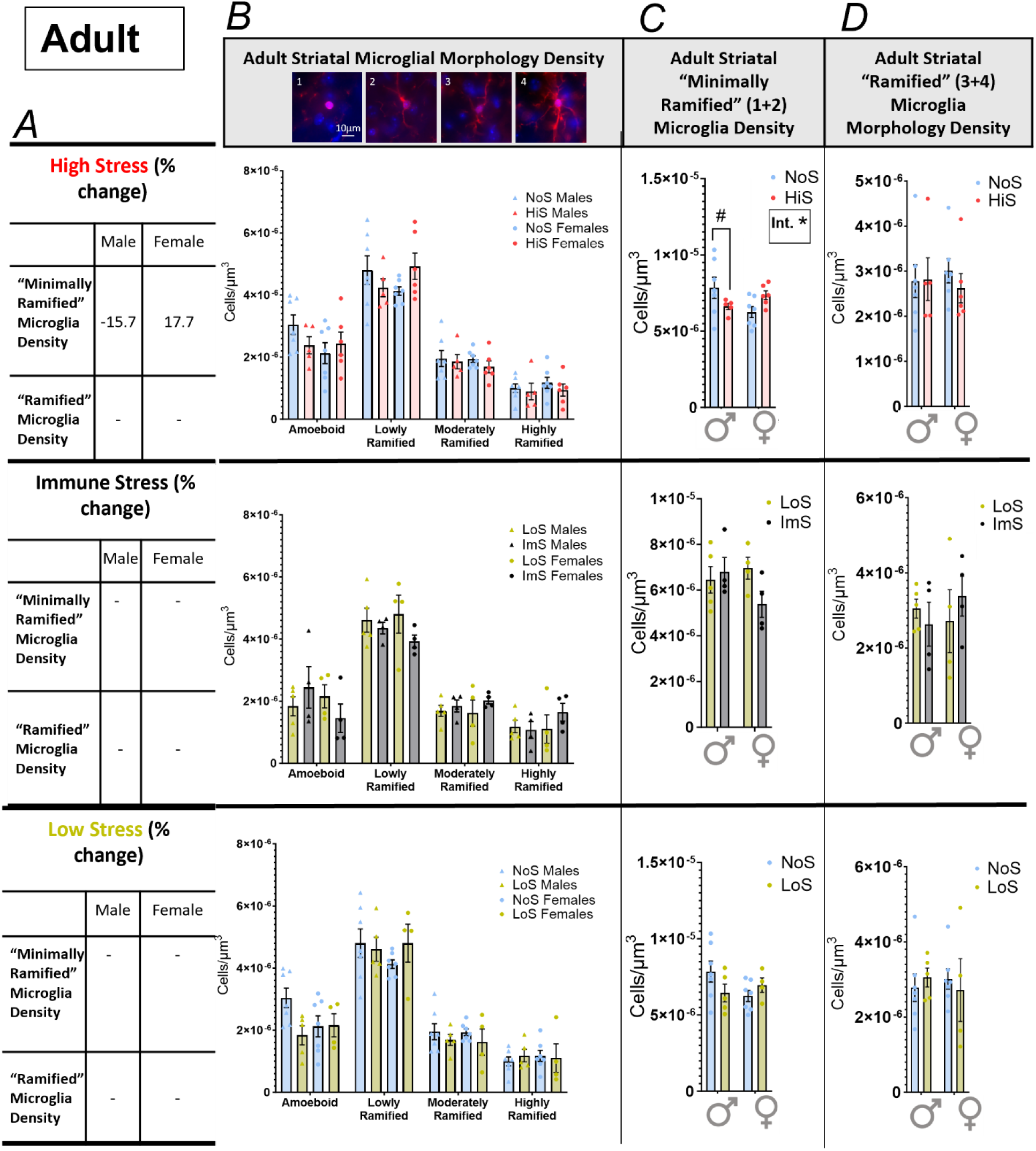
Adult microglial morphology changes due to stress. * *p* < 0.05, # *p* < 0.10, “Int.” = sex x stress interaction, NoS = No stress, LoS = Low stress, ImS = Immune stress, HiS = High stress. (A) Percent changes in each assessment and group for outcomes with *p* < 0.10 effects. (B) In adult offspring, there were no observed changes due to stress on microglial morphologies individually; 1 = amoeboid, 2 = lowly ramified, 3 = moderately ramified, 4 = highly ramified, Blue = DAPI, Red = IBA1. (C) When combining the two categories with low ramification, a significant sex x stress interaction was observed due to high stress. (D) No differences were observed after combining the two more ramified categories.

### Adolescent striatal-dependent behavior

To understand the developmental origins of striatal-dependent behavioral effects in adult offspring, we also assessed behavior at the adolescent timepoint (Fig 4*A*). HiS again had the most impacts across behavior, with minimal trend effects of ImS and LoS at this earlier stage. HiS was also the only model which reduced adolescent offspring weight (Supplemental Table S1) and also decreased locomotion in female offspring (Supplemental Fig S2*A*).

**Figure 4.**
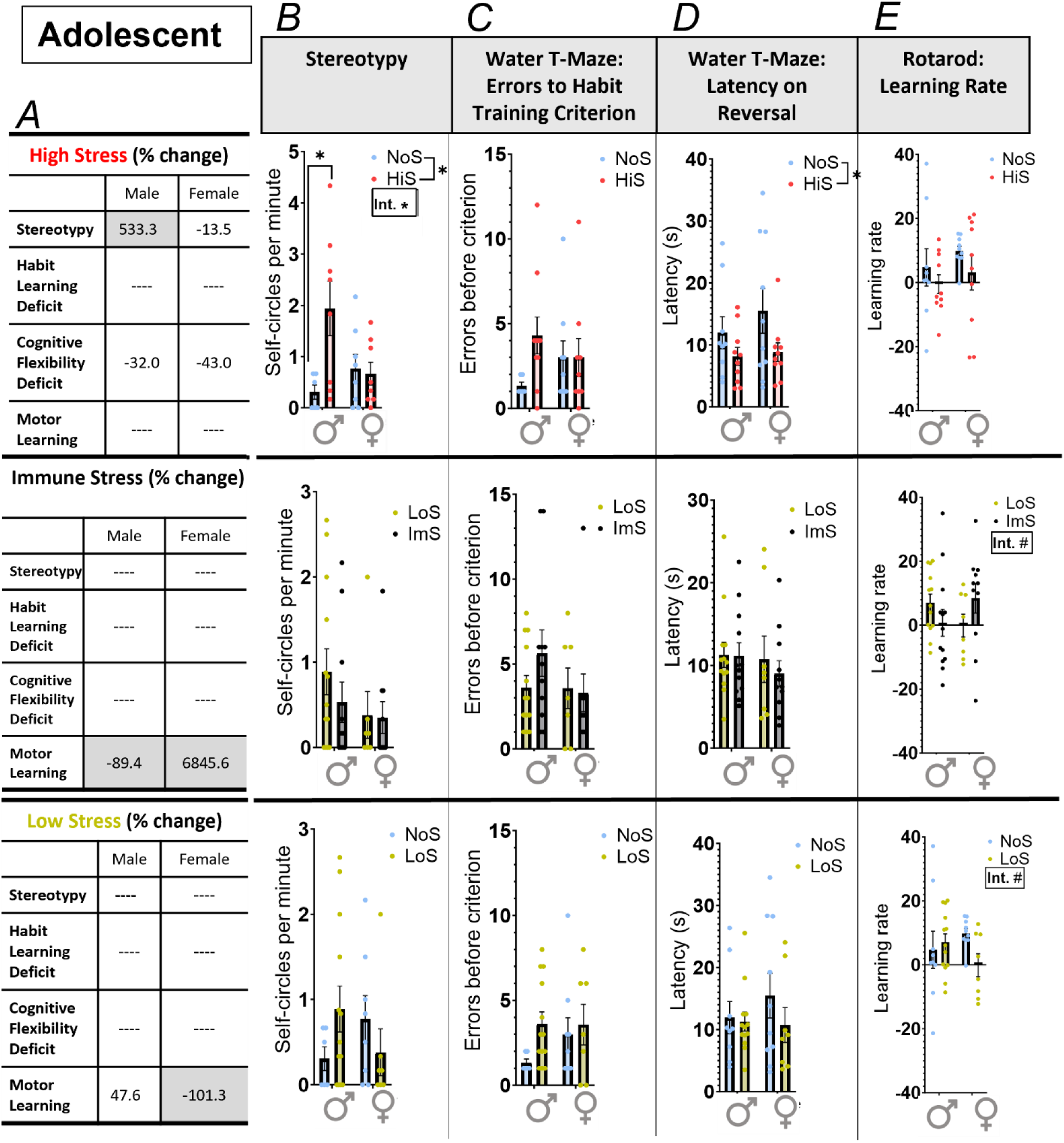
Juvenile striatal-dependent behaviors were largely unchanged. * *p* < 0.05, # *p* < 0.10, “Int.” = sex x stress interaction, NoS = No stress, LoS = Low stress, ImS = Immune stress, HiS = High stress. (A) Percent changes in each task and group. Cells that are shaded indicate a greater than 50% difference from control. (B) High stress led to an increase in stereotyped behavior, particularly in males. (C) No differences were observed in habit learning. (D) High prenatal stress led to a decrease in latency in the third reversal trial of the water T-maze, indicating a possible improvement in cognitive flexibility. (E) Motor learning was mostly unaffected by sex or stress condition, with a marginal interaction seen in the LoS vs ImS and NoS vs LoS comparisons.

**Figure 5.**
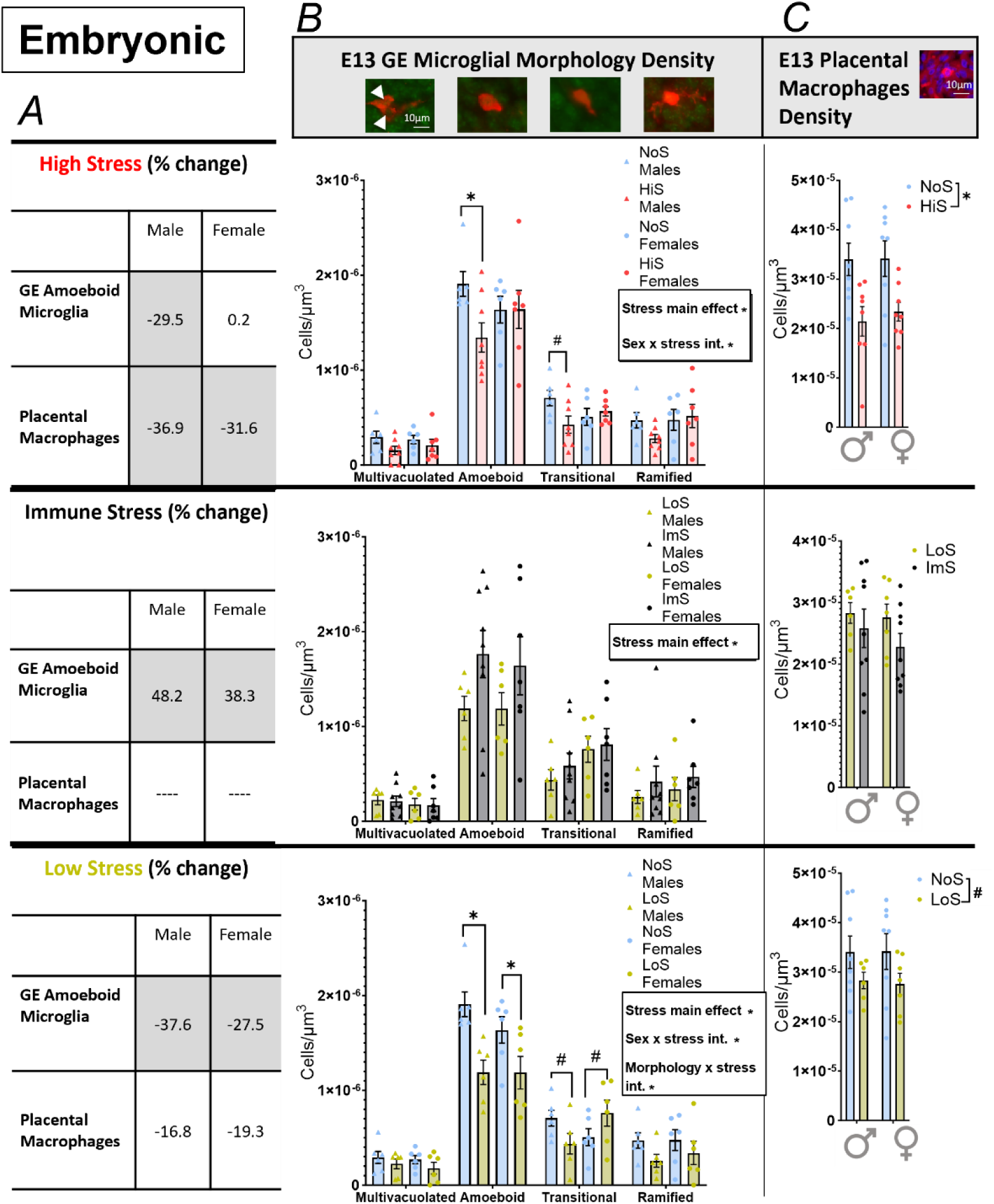
E13 changes due to stress. * *p* < 0.05, # *p* < 0.10, “Int.”= interaction. GE= ganglionic eminence, NoS = No stress, LoS = Low stress, ImS = Immune stress, HiS = High stress. (A) Percent changes in each assessment and group. Cells that are shaded indicate a greater that 25% difference from control. (B) In embryonic offspring, there were effects of all three stress groups on microglial morphology – with high and low leading to a decrease, particularly in amoeboid microglia, and immune stress leading to an increase in microglial density, from left to right: multivacuolated, amoeboid, transitional, ramified; Green = DAPI, Red = IBA1. (C) Placental macrophages were similarly decreased by high and low stress, Blue = DAPI, Red = IBA1.

HiS increased stereotyped behaviors in adolescent offspring, driven by a 533% higher level of self-circling in males (NoS vs HiS: F_1,26_ = 4.79, *p* = 0.038; sex x stress interaction: F_1,26_ = 6.18, *p* = 0.020, post hoc NoS vs HiS in males: *p* = 0.004, Fig 4*B*). This was particularly important given increased stereotyped behavior in HiS offspring even before amphetamine treatment (Supplemental Figure S2*C*). However, unlike in adult offspring, no effects of ImS were observed.

In the water-T maze, adolescent habit learning was not affected by any prenatal stress model unlike HiS and ImS effects on adults (Fig 4C). Also distinct from adults, HiS reduced latency to reversal learning in adolescent offspring (NoS vs HiS: F_1,35_ = 4.57, *p* = 0.040, Fig 4D), indicating possible enhancement in cognitive flexibility at this earlier time point.

Motor learning on the second day of the rotarod in adolescence was trend affected in sex-specific ways by ImS, but again with opposite effects as seen in adults (LoS vs ImS: sex x stress interaction: F_1,41_ = 3.47, *p* = 0.070, Fig 4*D*). Adolescent offspring exposed to LoS also had trend sex-specific changes in motor learning (NoS vs LoS: sex x stress interaction: F_1,36_ = 2.90, *p* = 0.097). Similar to adulthood, all adolescent offspring groups showed similar motor learning on day one (Supplemental Fig S2*F*).

Overall, adolescent behavior was not robustly altered with the exception of HiS impacts on male stereotyped behavior as also seen in adulthood.

### Early embryonic outcomes

To assess the origins of later behavioral and microglial changes, we turned our assessments to embryonic stages of development. First, both HiS and LoS decreased E13 body weight (Supplemental Table S1), suggesting that thresholds for effects at this early stage of development were easily exceeded.

Indeed, microglial density in the E13 primordial striatum (Fig 5*B*) - the ganglionic eminence - was affected by all models of stress (NoS vs HiS: F_1,92_ = 7.12, *p* = 0.009; LoS vs ImS: F_1,96_ = 5.40, *p* = 0.022; NoS vs LoS: F_1,80_ = 16.02, *p* = 0.0001). HiS and LoS decreased while ImS increased overall E13 microglial density (Fig 5*B*). Effects of HiS and LoS were sex-specific (sex-stress interactions: NoS vs HiS: F_1,92_ = 8.21, *p* = 0.005; NoS vs LoS: F_1,80_ = 4.01, *p* = 0.049) and affected some morphologies more than others (morphology-stress interaction: NoS vs LoS: F_3,80_ = 5.94, *p* = 0.001). HiS decreased E13 microglia only in males (post hoc: amoeboid *p* = 0.0002, transitional *p* = 0.061), similar to decreased minimally ramified microglia only in adult males. Despite no effect of LoS on adult striatal microglia, male and female E13 amoeboid microglia were decreased by LoS (post hoc: male *p* < 0.0001, female *p* = 0.004), while E13 transitional microglia were trend decreased in males and increased in females (post hoc: male *p* = 0.061, female *p* = 0.074). These robust effects occurred without changes to E13 ganglionic eminence volume or the E13 density of proliferating cells which would contribute to later growth (Supplemental Fig S3*A*,*B*).

The impact of both HiS and LoS at these early stages was also demonstrated by significant impacts on GAD67GFP+ cell migration out of primordial striatum (Supplemental Fig S3*C*).

We also assessed immune outcomes at the maternal-fetal gateway—the placenta. Both microglia and placental macrophages arise from a monocyte lineage (55) and may be affected with similar thresholds by prenatal stress. E13 placental macrophage density was decreased by HiS (NoS vs HiS: F_1,28_ = 14.99, *p* = 0.0006, Figure 5*C*) but also marginally by LoS (NoS vs LoS: F_1,25_ = 4.20, *p* = 0.051) across males and females. While ImS did not affect placental macrophage density, it did decrease E13 labyrinth zone area in males (Supplemental Fig S3*D*). HiS reduced labyrinth zone area in both males and females, and LoS did not affect the area of this zone (Supplemental Fig S3*D*). These results show that placental impacts occurred at this early stage across all stress models.

## Discussion

Consistent with our hypothesis, the low prenatal stress model was sufficient to affect early immune outcomes in embryonic brain and placenta, particularly in striatal microglia. However, only high and immune prenatal stress models led to persistent changes in adulthood striatal-dependent behavior or, for high stress only, adult striatal microglia. While impacts of the three prenatal stress models differed in their magnitude, direction, and effects on males and females, our results reveal that stress impacts on early outcomes appear greater, exceeding thresholds for effects more easily than for outcomes measured months after these prenatal manipulations (Fig 6).

**Figure 6.**
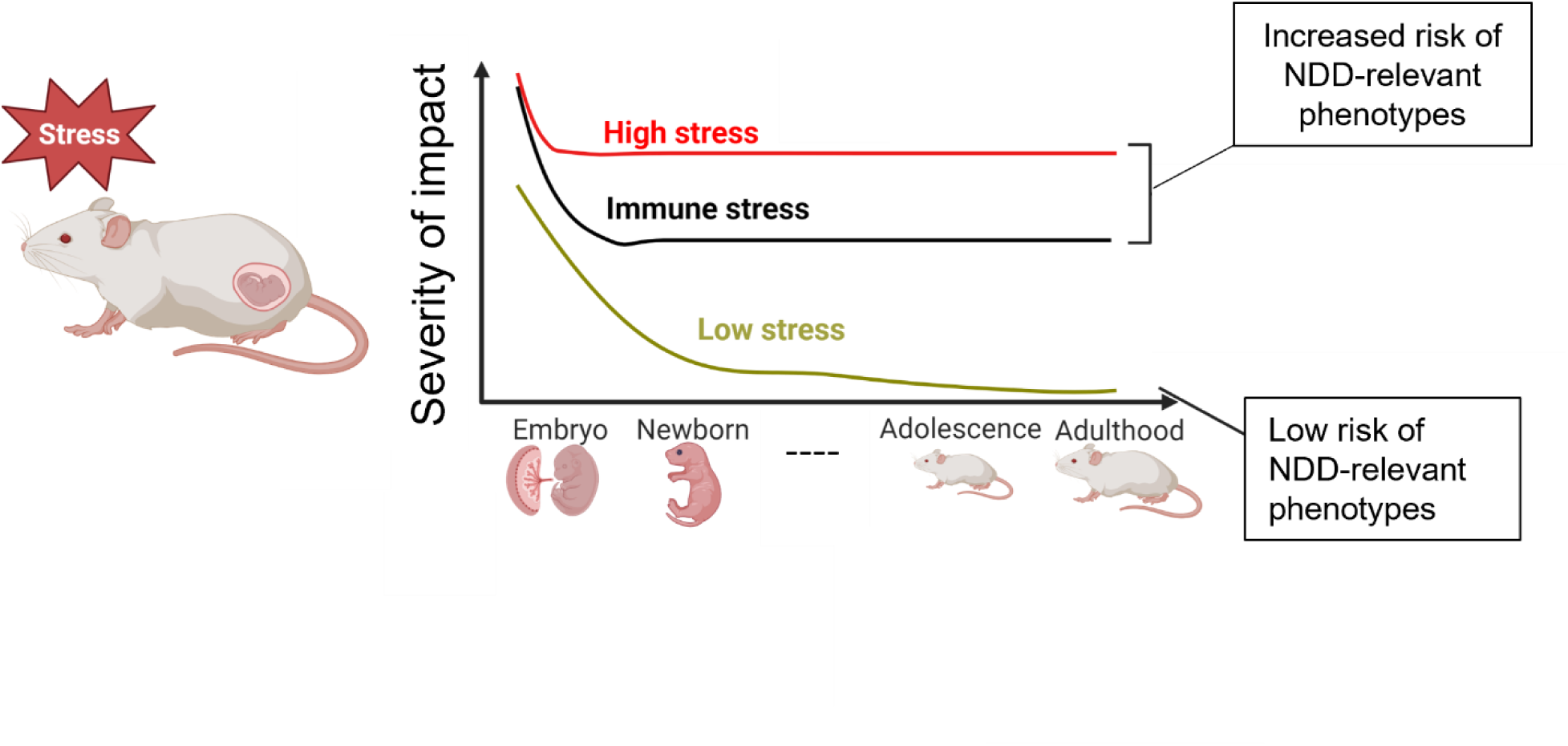
A proposed model of vulnerability in the context of age and stress severity. All stress models had early impacts, but long-term developmental outcomes were mostly induced by high stress. Immune stress also had some early and persistent effects like high stress but was not equivalent.

The impacts of prenatal stress on striatal-dependent behavior included increased stereotyped movements, quantified here as self-circling. Consistently across different stress types and in adolescence for high stress, male offspring stereotyped movements were more affected by prenatal stress.

Stereotyped movements are a feature across multiple neurodevelopmental disorders with links to prenatal stress and male predominance (56–58). Males were also more affected by the high stress model in adult striatal-dependent learning. Male-biased effects were not as clear for adult behavior changes due to immune stress or any stress effects on behavior in adolescence. This suggests that males may have different thresholds than females for prenatal stress effects on striatal function, especially those persisting across months.

Our findings also indicate that male striatal microglial morphology was altered due to high stress in both the adult and embryonic striatum consistent with their higher number of striatal-dependent behavioral disruptions. At the same time, striatal-dependent behavior and microglial outcomes were not consistently aligned within sex and stress type. This suggests that striatal microglia as a potential mechanism of prenatal stress programming may affect males and females differently or have different prominence. Previous findings have also suggested that striatal microglia may have a higher “threshold” of effect (59) than in other regions. A previous study found that maternal immune activation, a broad, one-time inflammatory challenge, decreased striatal microglial immune response in adulthood; this was similar to our results for high stress but not the more chronic, IL-6 immune stress used here (38). Our current work is one of the first to assess striatal microglia in the context of chronic prenatal stress. This striatal examination was all the more important due to these findings that impacts of prenatal stress differed for this specific region compared to other brain areas such as the cerebral cortex (19, 41, 54, 60).

NDDs associated with early-life risk factors and microglial changes include ASD and ADHD (61, 62)– however, much previous modeling of relevant ASD and ADHD mechanisms has lacked a striatal- centered approach. Our work converges with previous investigation of striatal microglia after maternal immune activation (MIA) (38). This showed reduced reactivity of microglia in the embryonic and adult striatum and striatal circuit changes which could be rescued by early replacement with control microglia (38). These findings add to other work implicating striatal microglia as critical modulators of neurodevelopment (40) and possible mediators of prenatal stress effects on striatal-dependent outcomes (63).

Outcomes assessed at the early time point, E13, revealed the lowest threshold for effects.

Embryonic microglia in the striatal primordium were decreased due to high and low stress, and increased due to immune stress. Interestingly, these early impacts recapitulate adult habit learning outcomes which showed opposite effects of high and immune stress. Despite this concordance, the low threshold of embryonic effects, evidenced by impacts of the minimal, low stress from saline injections alone, did not predict persistent behavioral effects in this model. There was some evidence that adult behavior and brain changes from this low level of prenatal stress were in similar direction to high stress effects but did not reach significance. This suggests the possibility of later compensation for early similar low and high stress impacts.

Because of previous findings implicating placental development and function in brain development (63, 64), we also assessed placental development as a function of prenatal stress severity. We found that the density of placental macrophages was decreased by both high and low stress – similar to decreased microglia seen in the embryonic brain. This suggests, as others have previously, that placental macrophages may act as a “window” to microglial development (55). Placental macrophages are critical for angiogenesis (65, 66), suggesting their importance in placental function and fetal development. However, immune stress increased placental macrophage density with opposite early embryonic brain microglia effects. Actions of IL-6 directly on placenta but more indirectly on the developing brain may underlie these differences.

Assessment at the adolescent intermediate time point also generally demonstrated increasing thresholds for more persistent effects over development. While adolescent offspring showed general (weight loss) effects of the high stress model, this and the immune stress model caused fewer behavioral alterations than in adults. Adolescence has previously been implicated as a particularly plastic time for brain development (67) and this may mask or compensate for effects of prenatal stress.

This investigation was not designed to evaluate other aspects of microglia function, which most certainly extends beyond their morphology (68). However, the evaluation of morphology across different developmental time points was revealing and the focus on the striatal microglia revealed substantially different impacts of prenatal stress than those seen in other brain regions. In addition, impacts from a low stress model used here and elsewhere often as a control condition for inflammatory exposure models (maternal vehicle injections) were distinct from non-manipulated controls in this work. Such differences have rarely been considered. However, this work had other limitations including behavioral tasks which involve striatum as well as other brain regions and the use of an outbred mouse line which may introduce variability.

## Conclusion

Overall, the current work indicates that age, sex, and severity all frame how prenatal stress affects brain and behavioral development. These findings support striatal microglia as a candidate mechanism for prenatal stress effects and suggest placental response may be a mediating factor. The interplay between microglia and striatal development as a function of stress may explain why some striatal-dependent behaviors were altered more than others. This work is critical to the future understanding of the early origins of NDDs, with the goal of tailoring individual circumstances with individual therapies. More work on striatal development is critical to improving mechanistic understanding of early risks for NDDs.

## Author Contributions

SVM and HES conceived the concept of the current work. SVM and MME designed the experiments with significant input from HES. SVM, MME, MD, SK, JLI, RME, SLA, and VCR collected data. SVM, MME, and MD analyzed data. SVM, MD, and HES wrote the manuscript, which all authors reviewed and/or edited.

