## Supplemental Information for "Threshold effects of prenatal stress on striatal microglia and relevant behaviors"

Supplemental Table 1.
Body weight data show a lower threshold for effect due to age and severity of stress, with high stress having a more long-lasting impact than immune stress and low stress. * $p < 0.05$ significantly different from the same-sex NoS group (HiS and LoS) or the same-sex LoS group (ImS). Mean (± standard error of the mean). P21 or P0 = postnatal day 21 or 0, E13 = embryonic day 13. NoS = No stress, LoS = Low stress, ImS = Immune stress, HiS = High stress.

| | Adult Males | Adult Females | P21 Males | P21 Females | P0 Sexes Combined | E13 Males | E13 Females |
|---|---|---|---|---|---|---|---|
| HiS | 41.88 (±1.631) | 33.87 (±1.240) | 15.63* (±0.391) | 15.30* (±0.298) | 1.58* (±0.045) | 0.185* (±0.004) | 0.177* (±0.004) |
| ImS | 42.59 (±1.083) | 33.85 (±0.993) | 16.29 (±0.392) | 15.30 (±0.393) | 1.63 (±0.040) | 0.178 (±0.004) | 0.176 (±0.006) |
| LoS | 42.02 (±1.262) | 33.78 (±1.341) | 16.79 (±0.473) | 15.84 (±0.464) | 1.69 (±0.043) | 0.176* (±0.008) | 0.171* (±0.006) |
| NoS | 43.32 (±0.923) | 33.98 (±0.830) | 16.97 (±0.202) | 15.62 (±0.171) | 1.81 (±0.045) | 0.204 (±0.005) | 0.191 (±0.003) |

Supplemental figure 1.

Adult behavioral and brain data. # p < 0.10, NoS = No stress, LoS = Low stress, ImS = Immune stress, HiS = High stress. (A) No group differences were observed in time mobile in the open field task. (B) No group differences were observed in speed while mobile in the open field task. (C) Prior to amphetamine injection, no group differences were observed in self-circling behavior. (D) In the water T-maze, a marginal group difference in the latency of the first reversal trial was observed between the LoS and ImS groups, with ImS offspring being faster. (E) No group differences were observed in the latency of the second reversal trial in the water T-maze. (F) No group differences were observed in learning rate of the first day of rotarod training. (G) No differences were observed in adult striatal volume.

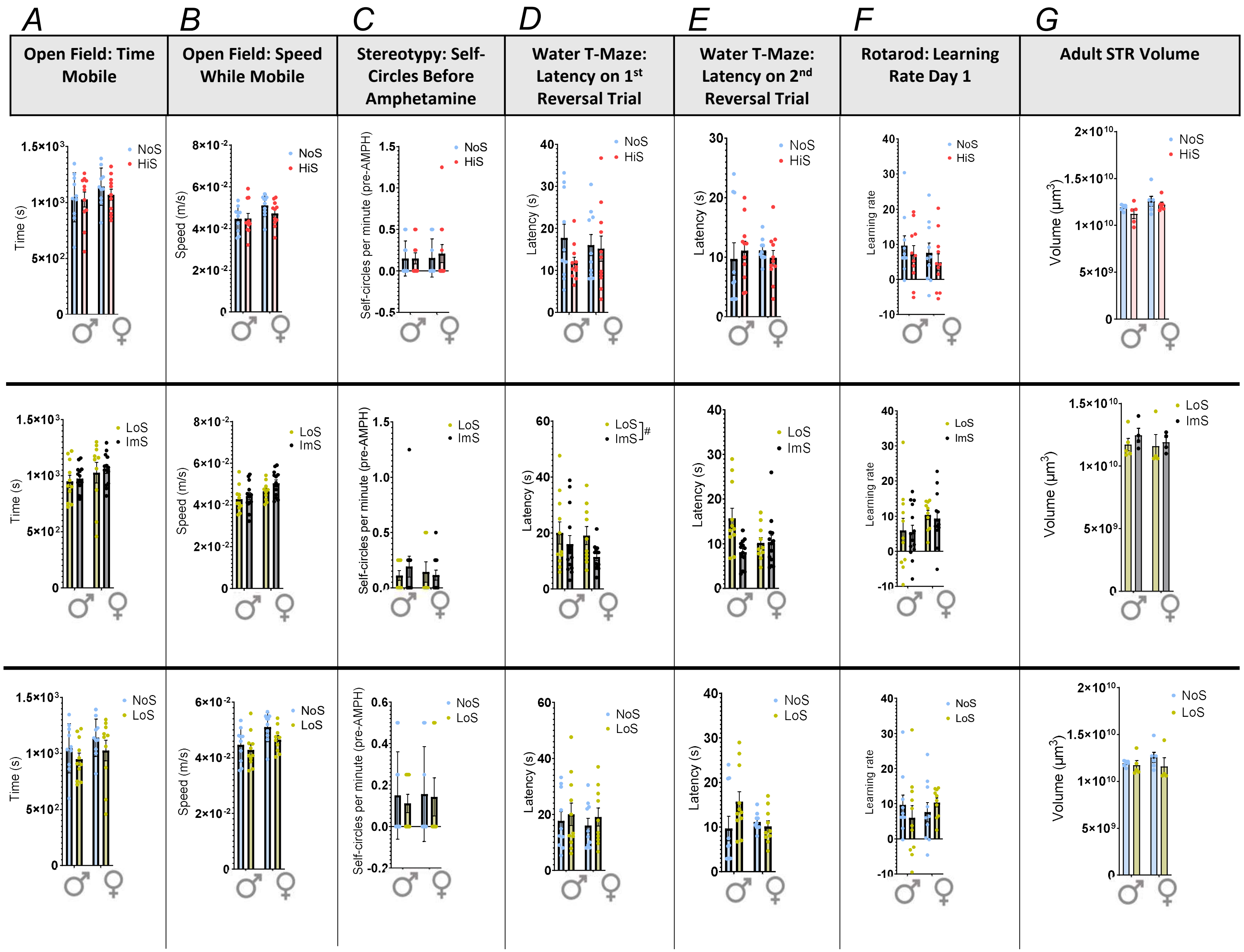

Supplemental figure 2.

Adolescent behavioral data. * $p < 0.05$, # $p < 0.10$, NoS = No stress, LoS = Low stress, ImS = Immune stress, HiS = High stress. (A) In the open field task, a significant sex x stress interaction was observed between the NoS and HiS groups, with females spending trending less time mobile. (B) No group differences were observed in speed while mobile in the open field task. (C) Prior to amphetamine injection, a significant increase in self-circling behavior was observed in juvenile high stress offspring, possibly indicating a baseline increase in stereotyped behaviors. This group difference perseveres after amphetamine injection, particularly in males, but self-circling behavior is notably increased by amphetamine. (D) No group differences were observed in the latency of the first reversal trial in the water T-maze. (E) In the latency of the second reversal trial, a significant sex x stress interaction was observed in the HiS group. (F) No group differences were observed in learning rate of the first day of rotarod training.

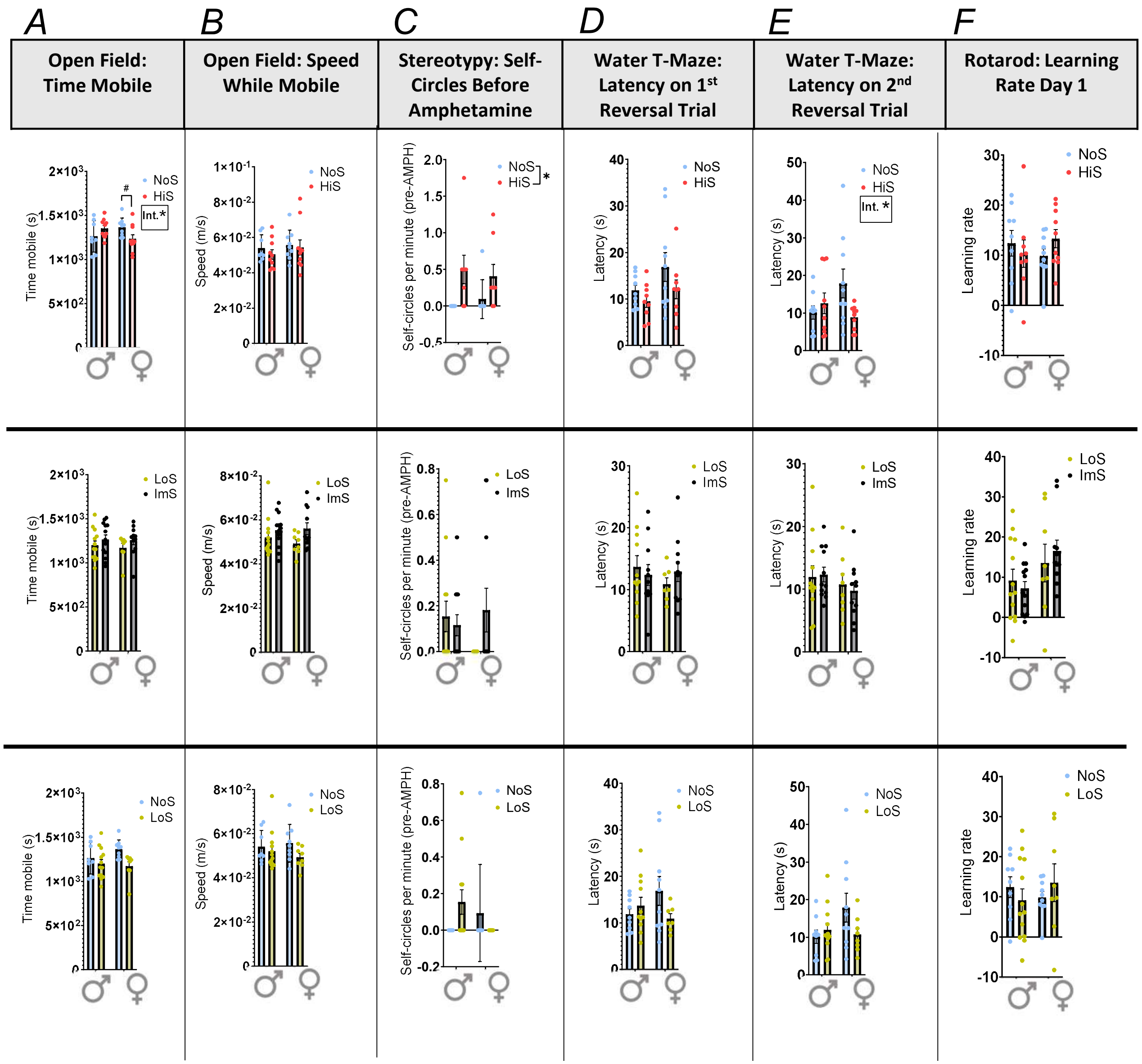

Supplemental figure 3.

Embryonic brain and placenta measures. * p < 0.05, # p < 0.10, “Int.” = interaction, GE = ganglionic eminence, NoS = No stress, LoS = Low stress, ImS = Immune stress, HiS = High stress. (A) E13 GE volume was unchanged by stress. (B) Cell division was not altered by stress. (C) High and low stress led to a significant delay in GABAergic cell migration across the cortex. (D) High and immune stress led to area decreases in the labyrinth zone of the placenta – in both sexes due to high stress, and in males only due to immune stress. (E) No differences were observed in placental junctional zone area.

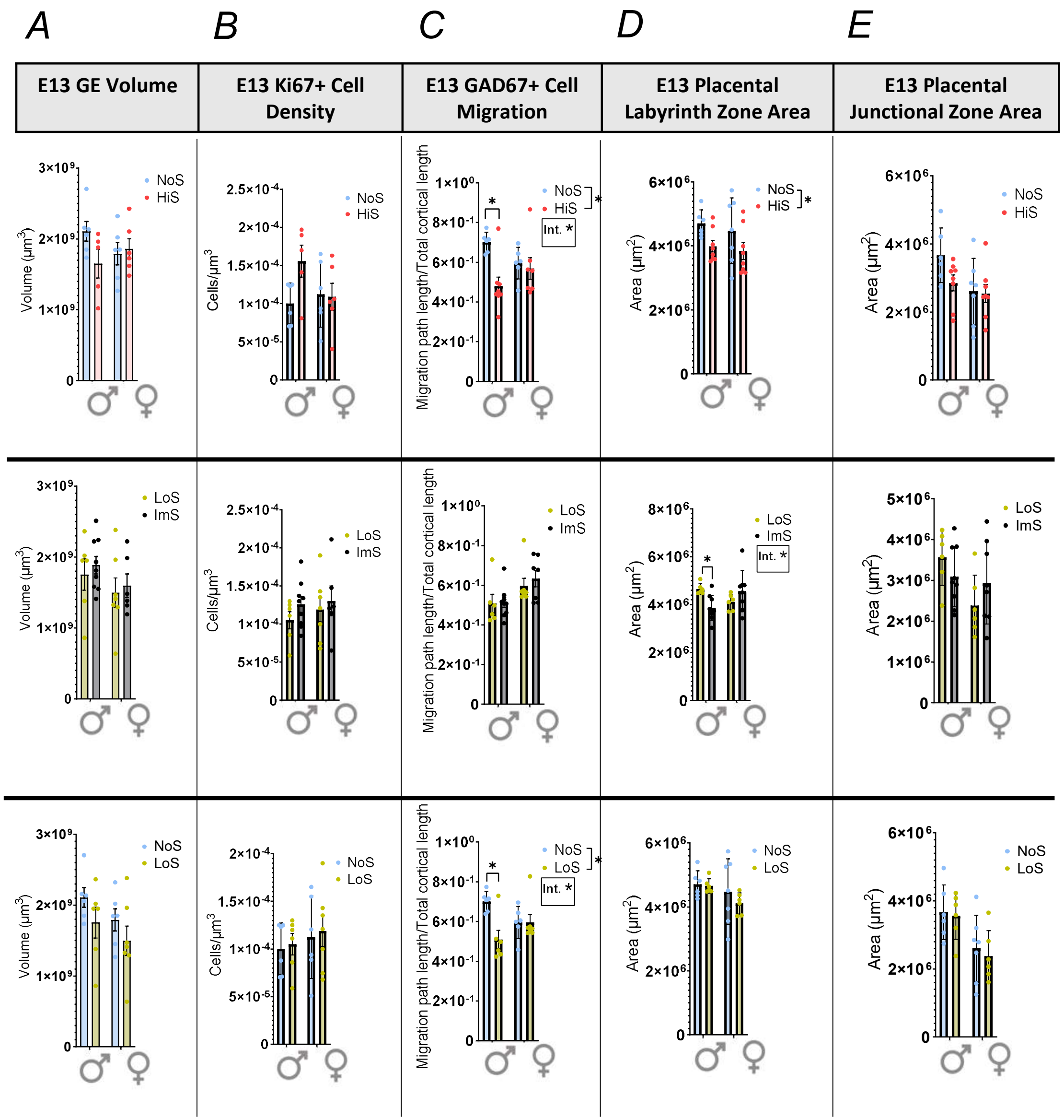
